# COVID-19 Infection and Transmission Includes Complex Sequence Diversity

**DOI:** 10.1101/2022.04.18.488717

**Authors:** Ernest R. Chan, Lucas D. Jones, Marlin Linger, Jeffrey D. Kovach, Maria M. Torres-Teran, Audric Wertz, Curtis J. Donskey, Peter A. Zimmerman

## Abstract

SARS-CoV-2 whole genome sequencing has played an important role in documenting the emergence of polymorphisms in the viral genome and its continuing evolution during the COVID-19 pandemic. Here we present data from over 360 patients to characterize the complex sequence diversity of individual infections identified during multiple variant surges (e.g., Alpha and Delta; requiring ≥ 80% genome coverage and ≥100X read depth). Across our survey, we observed significantly increasing SARS-CoV-2 sequence diversity during the pandemic and frequent occurrence of multiple biallelic sequence polymorphisms in all infections. This sequence polymorphism shows that SARS-CoV-2 infections are heterogeneous mixtures. Convention for reporting microbial pathogens guides investigators to report a majority consensus sequence. In our study, we found that this approach would under-report at least 79% of the observed sequence variation. As we find that this sequence heterogeneity is efficiently transmitted from donors to recipients, our findings illustrate that infection complexity must be monitored and reported more completely to understand SARS-CoV-2 infection and transmission dynamics involving both immunocompetent and immunocompromised patients. Many of the nucleotide changes that would not be reported in a majority consensus sequence have now been observed as lineage defining SNPs in Omicron BA.1 and/or BA.2 variants. This suggests that minority alleles in earlier SARS-CoV-2 infections may play an important role in the continuing evolution of new variants of concern.

**AUTHOR SUMMARY:** Evolution of the virus causing COVID-19 (SARS-CoV-2) has been associated with significant transmission surges. With evolution of SARS-CoV-2, evidence has accumulated regarding increased transmissibility of lineages, varying severity of illness, evasion of vaccines and diagnostic tests. Continuous tracking of SARS-CoV-2 lineage evolution distills very large and complex viral sequence data sets down to consensus sequences that report the majority nucleotide at each of over 29,000 positions in the SARS-CoV-2 genome. We observe that this eliminates considerable sequence variation and leads to a significant underestimation of SARS-CoV-2 infection diversity and transmission complexity. Additionally, concentration on the majority consensus sequence diverts attention from genetic variation that may contribute significantly to the continuing evolution of the COVID-19 pandemic.

## INTRODUCTION

Early investigation into the outbreak of the novel pneumonia of unknown cause, from the city of Wuhan in late December 2019, quickly identified the genomic sequence [1, 2] associated with a coronavirus, closely related to the SARS-CoV [3, 4]. The virus, SARS-CoV-2, is now understood to be the causative agent of COVID-19. Sequence variation across the ∼30kb positive-strand RNA SARS-CoV-2 genome was reported in the earliest studies, including single nucleotide polymorphisms (SNP), insertions and deletions (indels) [1, 5, 6]. Significant efforts have since focused on characterizing sequence variation and documenting the evolution of SARS-CoV-2 strains during the pandemic through over 1.8 million sequences submitted from 172 countries as of April 2021 [7]. Extensive effort [8–11] has and continues to perform phylogenetic assessments across the ongoing evolution of SARS-CoV-2 populations around the world. This effort has captured and presented evidence of the continuous emergence of the major SARS-CoV-2 lineages sharing sequence variations across the global landscape during the pandemic. Many notable lineages including, B.1.1.7 (Alpha), B.1.351 (Beta), P.1 (Gamma), B.1.617.2 (Delta) and B.1.1.529 (Omicron) have been associated with important surges in COVID-19 transmission and on-going phases of the pandemic [9, 12]. With the surge of each new variant, concern naturally arises (and evidence accumulates) about strain-specific transmission and virulence, as well as the effectiveness of COVID-19 diagnostics, therapeutics, and vaccines.

From earlier studies of the SARS-CoV-1 virus, it was suggested that the error-correcting capacity of the RNA-dependent RNA polymerase (RdRP) conferred by nonstructural protein 14 (NSP14) [13, 14] would contribute to replication error rates significantly lower than other RNA viruses (e.g. HIV-1 and influenza virus) [15, 16]. Studies comparing SARS-CoV-1 and SARS-CoV-2 estimate mutation rates that range from 1–3 × 10^−3^ substitutions/site/year [8, 17–19], respectively (extrapolating to 30-90 mutations/genome/year) are consistent with this prediction. Data provided through the Global Initiative on Sharing All Influenza Data (<u>GISAID</u>), and NextStrain provides a continuously updating report on SARS-CoV-2 lineage classification[20, 21] and illustrates that an apparently low mutation rate is no guarantee that the virus will exhibit limited capacity for variation, particularly once it has become so widely dispersed through millions of daily infections [6, 22, 23].

Data analyzed for lineage tracking relies on extensive curation and validation of SARS-CoV-2 genomic data. This includes minimization and resolution of ambiguous sequence data. In fact, most sequence data used to monitor the COVID-19 pandemic distills the whole genome sequence generated from an infection sample, to a single consensus sequence (the majority nucleotide present at each genomic position) [24–27] to national and international databases [28, 29]. Out of the wide range of SARS-CoV-2 genomics studies, sequence variation within infected individuals has been described in relatively few studies as intra-host single nucleotide variants (iSNVs) [30–38]. Outcomes of these studies appear to be mixed in their assessments of individual infection diversity and transmission outcomes but inevitably reveal and acknowledge that these variations within a single infection exists.

Here we evaluate SARS-CoV-2 whole genome data from a sample of patients, administrative and medical staff experiencing COVID-19 in the VA Northeast Ohio Healthcare System. We have applied stringent inclusion criteria based on SARS-CoV-2 genome coverage and read-depth to normalize comparisons across sequences derived independently from individual patient isolates as well as patients who have had direct prolonged contact with other infected individuals. We examine multiple perspectives, including details of individual infection diversity and transmission outcomes which are critical for the successful management of the COVID-19 pandemic.

## RESULTS

### SARS-CoV-2 Genome Sequence Coverage and Detection of Genetic Variation

A total of 364 SARS-CoV-2 full genome sequences were analyzed in comparison to the Wuhan reference genome (GenBank <u>NC_045512</u>). A total of 254 samples from patients (N=179) and healthcare personnel (N=75) with COVID-19 were collected to (a) perform high-resolution contact tracing of COVID-19 infections and (b) gain insight into SARS-CoV-2 transmission dynamics between donor and recipient individuals [36–38]. Because of the infection control and outbreak monitoring nature of this work, the individuals represented a wide demographic range, including gender, age, race, and health status. Of 175 patients with available clinical information, the mean age was 61.4 years (range, 24 to 98), 31 (18%) were asymptomatic, 58 (33%) were hospitalized for COVID-19 or non-COVID-19 conditions, 32 (18%) had severe COVID-19, and 11 (6%) died due to complications of COVID-19. Twenty-four (14%) patients with COVID-19 were moderately or severely immunocompromised, including 17 with active treatment for malignancy with chemotherapy or severely immunosuppressive medications, 2 with advanced or untreated HIV infection, 1 with an organ transplant requiring immunosuppressive therapy, and 4 with other chronic medical conditions requiring immunosuppressive therapy. All patient samples have been analyzed as part of an effort to assess the extent of sequence variability within SARS-CoV-2 infections across different surges associated with variants of interest and concern from June 2020 to October 2021. An additional 110 samples from the Sequence Read Archive (SRA) have been analyzed for assessment of diversity outside the local region.

Each sample was initially evaluated by RT-PCR to approximate levels of infection that would enable high-resolution evaluation of individual infection complexities. The Ct values for samples that resulted in SARS-CoV-2 full genome coverage (≥80% of the genome covered at ≥100X read-depth) were significantly lower (Average Ct value of 27.96 [n= 138]), implying higher viral RNA copy number than samples that did not meet this coverage threshold (Average Ct value of 35.85 [n=103]) (**Fig. 1a**; Mann-Whitney U= 1,971.5, P<0.0001). Coverage across the SARS-CoV-2 genome for samples with sequence generated above (black line) and below (gray line) the coverage threshold is summarized in **Fig. 1b** and direct correlation between Ct and average coverage for each individual sample is provided in **Fig. 1c**. Among the 140 (two samples did not have RT-PCR data) sequences meeting our technical thresholds, the average coverage was 1,081X (range 223 to 2,235X) with an average of 279,408 uniquely mapped reads per sample (min=63,296, max=2,680,239). Further examination of our data showed that the ability to detect SNPs was not influenced by Ct or average coverage if a minimum coverage was reached (**Figs. 1d** **and 1e**). Samples that pass our inclusion criteria, ≥80% of the genome covered at ≥100X read depth, showed comparable number of SNPs defined by alternate allele frequency (AAF) > 5% (filled black circles) or SNPs defined by AAF > 50% (filled diamonds), regardless of Ct value. As expected, fewer SNPs were detected in samples that did not meet our inclusion (open circles and diamonds). With read depth of ≥100X, we observed consistency in detecting expected sequence variations within our sampled population and across genomic sequences reported into the international databases. From these analyses we established confidence in detecting minor allele at frequencies ≥5% (whether reference or alternate allele). For the 114 samples not meeting our criteria for inclusion, the average coverage was 71X (range 0 to 1,281X) and these samples were removed from subsequent analyses. A comparison of the genome sequence data across the total sampling period of our survey showed a significant increase in the number SNPs detected over time (**Fig. 1f** **and Table 1**) with average coverage remaining consistent through this period; approximately doubling (30 to 65 AAF ≥5%) or tripling (10 to 35 AAF ≥50%) from April 2020 to August 2021. The two samples demonstrating substantially higher SNP counts (**Figs. 1d-1f red and green arrows**) will be covered in their epidemiological context below.

**Fig. 1.**
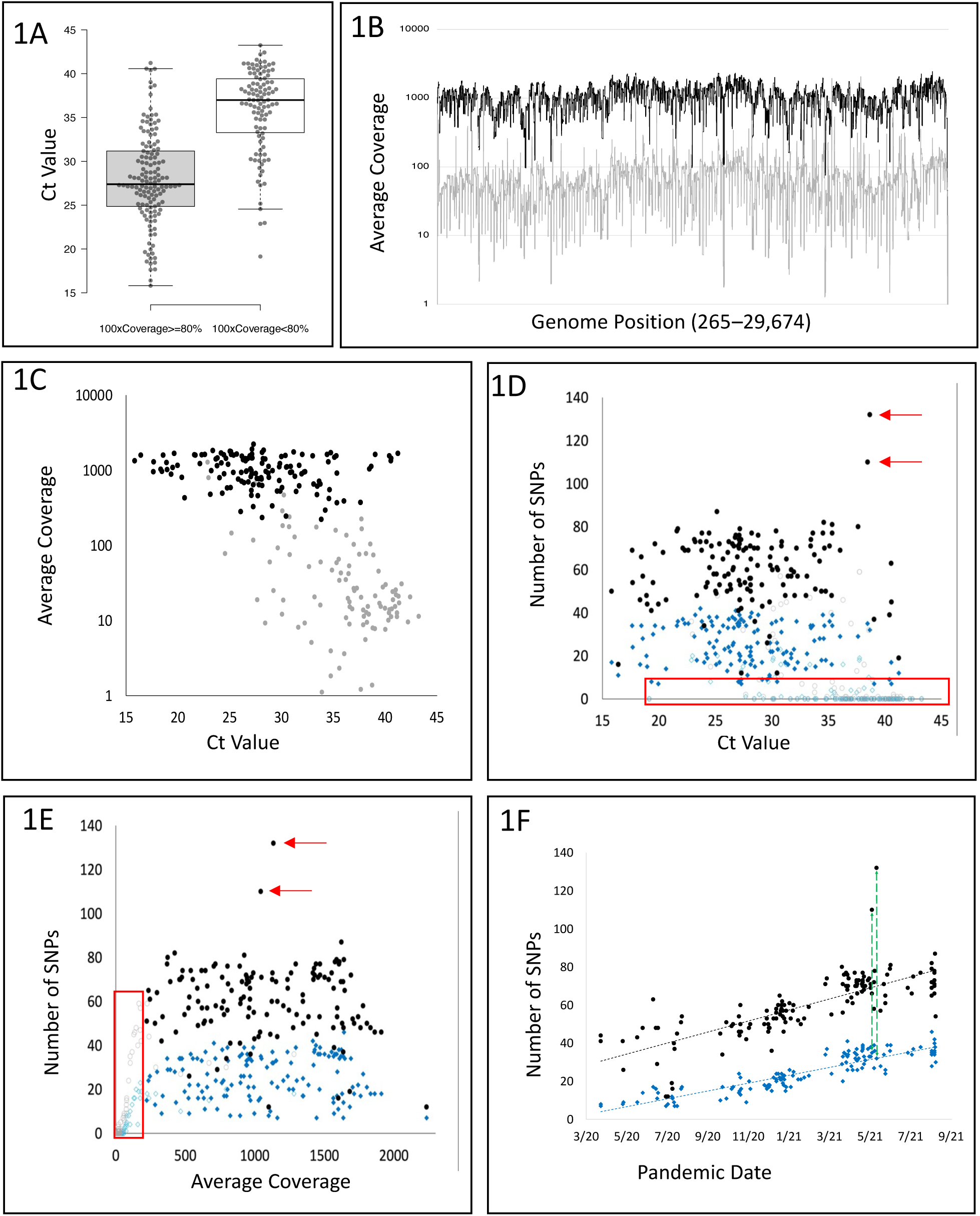
Assessment SARS-CoV-2 Amplicon Library Preparation and Genome Coverage. **a** RT-PCR cycle thresholds predicting SARS-CoV-2 sequence read quality. Comparison of RT-PCR cycle thresholds for samples according to technical breadth and depth thresholds ((≥80% of SARS-CoV-2 genome covered at ≥100X read-depth; met thresholds, n= 140; did not meet thresholds n= 114). **b** SARS-CoV-2 full genome sequence depth comparisons. Among 140 sequences meeting our technical thresholds, the average coverage was 1,081 X (range 223 to 2,235 X); for the 114 sequences not meeting our technical thresholds, the average coverage was 71 X (range 0 to 1,281 X). Among all samples, average uniquely mapped read count was 292,430 reads (min=21, max=2,680,239). Among samples that pass >80% 100X coverage, the average uniquely mapped read count was 279,408 reads (min=63,296, max=2,680,239). Among samples that did not pass our cutoff, the average uniquely mapped read count was 175,895 reads (min=21, max=1,283,414). **c** Correlation between RT-PCR Ct value and Average viral genome coverage; black circles identify samples that met the coverage and read-depth thresholds, and gray circles identify samples that did not meet these thresholds. **d** Correlation between detection of alternate alleles and RT-PCR Ct value where alternate allele frequency (AAF) was >5% (black circles >80%100x and grey circles <80%100x), or where AAF was >50% (blue diamond >80% 100x, blue triangles <80% 100x). **e** Correlation between detection of alternate alleles and average SARS-CoV-2 genome coverage where alternate allele frequency (AAF) was >5% (black circles >80%100x and grey circles <80%100x), or where AAF was >50% (blue diamond >80% 100x, blue triangles <80% 100x). **f** Correlation between the number of alternate alleles identified in the 140 >80% 100x samples and their collection dates - AAF >5% (black circles), or AAF >50% (blue diamonds). Significant correlation was observed between an increase in AAFs during the sample collection time-period (AAF >5%, Pearson R = 0.76112; AAF >50%, Pearson R = 0.89253; both P-values <0.0001). Red arrows in **d-e** identify samples that carried significantly more sequence variation compared to expectation (Surv20 – 110 SNPs; Surv42 – 132 SNPs). Green arrows in **f** link the position of the two samples with additional sequence variation at the AAF >50% and AAF >5% thresholds.

**Table 1.** Assessment of SNVs at Time Quartiles of the COVID-19 Pandemic.

| **Table 1. Assessment of SNVs at Time Quartiles of the COVID-19 Pandemic** |  |  |  |  |  |  |
| --- | --- | --- | --- | --- | --- | --- |
| Quartile | SNP Avg<br>(AAF>5%) | Min (>5%) | Max (>5%) | SNP Avg<br>(AAF>50%) | Min (>50%) | Max (>50%) |
| 8/5/20 | 37.74 | 12 | 63 | 10.89 | 7 | 17 |
| 12/9/20 | 48.68 | 34 | 60 | 16.16 | 9 | 24 |
| 4/14/21 | 59.91 | 36 | 80 | 24.45 | 15 | 41 |
| 8/18/21 | 72.33 | 40 | 132 | 33.76 | 8 | 46 |

### Consensus Sequences and SARS-CoV-2 Strain Polymorphism

Enumeration of SNPs across the SARS-CoV-2 genes and a full listing of all SNPs observed in our study are presented in **Supplemental Table 1** (SNPs observed in ≥10 infections) and **Supplemental Table 2** (all SNPs), respectively. Alignment of the 140 SARS-CoV-2 consensus sequences (**Supplemental Table 3**) that met the coverage thresholds, revealed a total of 1,096 SNV positions. Among these polymorphisms, 406 SNPs were observed in multiple infection samples.

The variant positions across the SARS-CoV-2 genomic sequences detected with allele frequencies >5% reveal extensive biallelic polymorphism throughout our results (**Fig. 2** **and** **Fig. 3**). We first examine these data by assessing alternate vs reference allele frequency proportions (AAF:RAF) for the 75 SNV positions that were shared most commonly (in ≥10 samples) (**Fig. 2**). The red dashed line in the figure at 50% allele frequency is the commonly accepted cutoff for variant detection indicating major allele. Only those alternate allelic SNPs with frequencies above 50% would be reported as part of the consensus sequence characterizing the virus present in the infection sample. Following this convention, 30 alternate allelic positions observed in our surveyed population would not have been reported in any of the infection-specific consensus sequences, totaling 2,865 SNP events. Among the 45 SNPs where AAFs did rise above the 50% threshold, the alternate allele would be reported in 1,143 occurrences (AAF >50%) and not reported in 1,553 (AAF 5-50%). Following the consensus sequence reporting convention across the 75 positions monitored, most of the SARS-CoV-2 variations (4,418 SNP out of 5,561, 79%) with alternate allele frequencies between 5-50% would not have been reported across the infections studied. These findings shown previously in **Figs. 1d-1f**, predicting lower levels of polymorphism at a 50% vs 5% threshold, are consistent with this analysis.

**Fig. 2.**
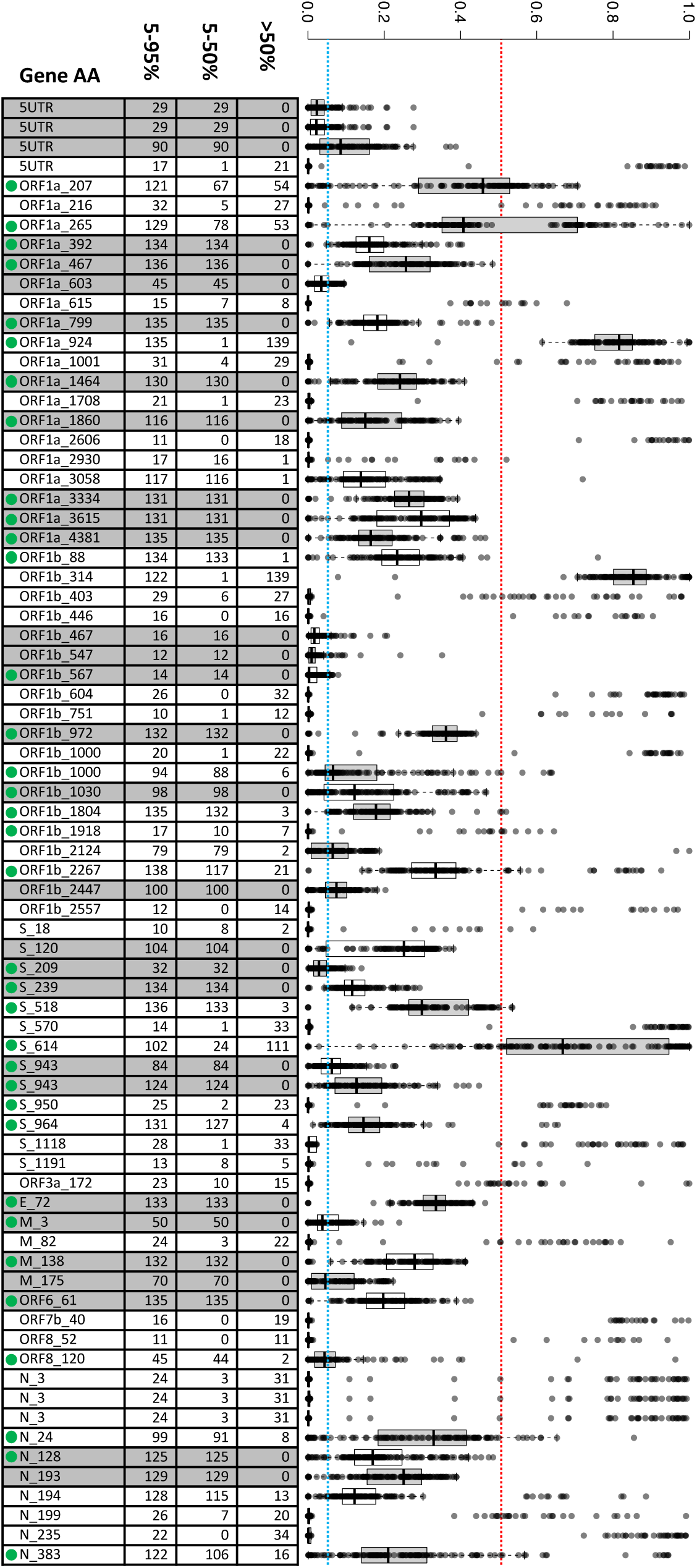
Assessment of Alternate and Reference Allele proportions (AAF:RAF) Across 140 Infections. Box and whisker plots illustrate the spread of RAF:AAF proportions for each mutation (median (bar); 25^th^ and 75^th^ percentiles (bottom and top of box, respectively); 10^th^ and 90^th^ percentiles (bottom and top whiskers, respectively)). The red dashed line at 50% identifies the demarcation at which the alternate allele is recognized and reported as the conventional consensus nucleotide that characterizes the infecting virus. These 75 mutations were chosen for further analysis because they all showed evidence of biallelic states at greater than the 5% background threshold and were shared in more than 10 infections (blue dashed line). Among these varying positions, Illumina sequence reads for 30 mutations did not reach an AAF:RAF >50% for any of the 140 infections; 45 positions exceeded the RAF:AAF 50% threshold for a portion of the 140 infections and would therefore have contributed to characterization of the infecting viral strain. The accompanying table enumerates the number of samples with AAF between 5 and 95%, <50% and >50%. Overall, among the 5,390 alternate alleles detected at a frequency >5%, just 975 (18%) were observed at >50% frequency necessary to be reported in consensus sequence. Nucleotide positions characterized by biallelic iSNVs in our VA study population and the samples queried in the SAR (Supplemental Fig. 1) are designated by a green dot.

**Fig. 3.**
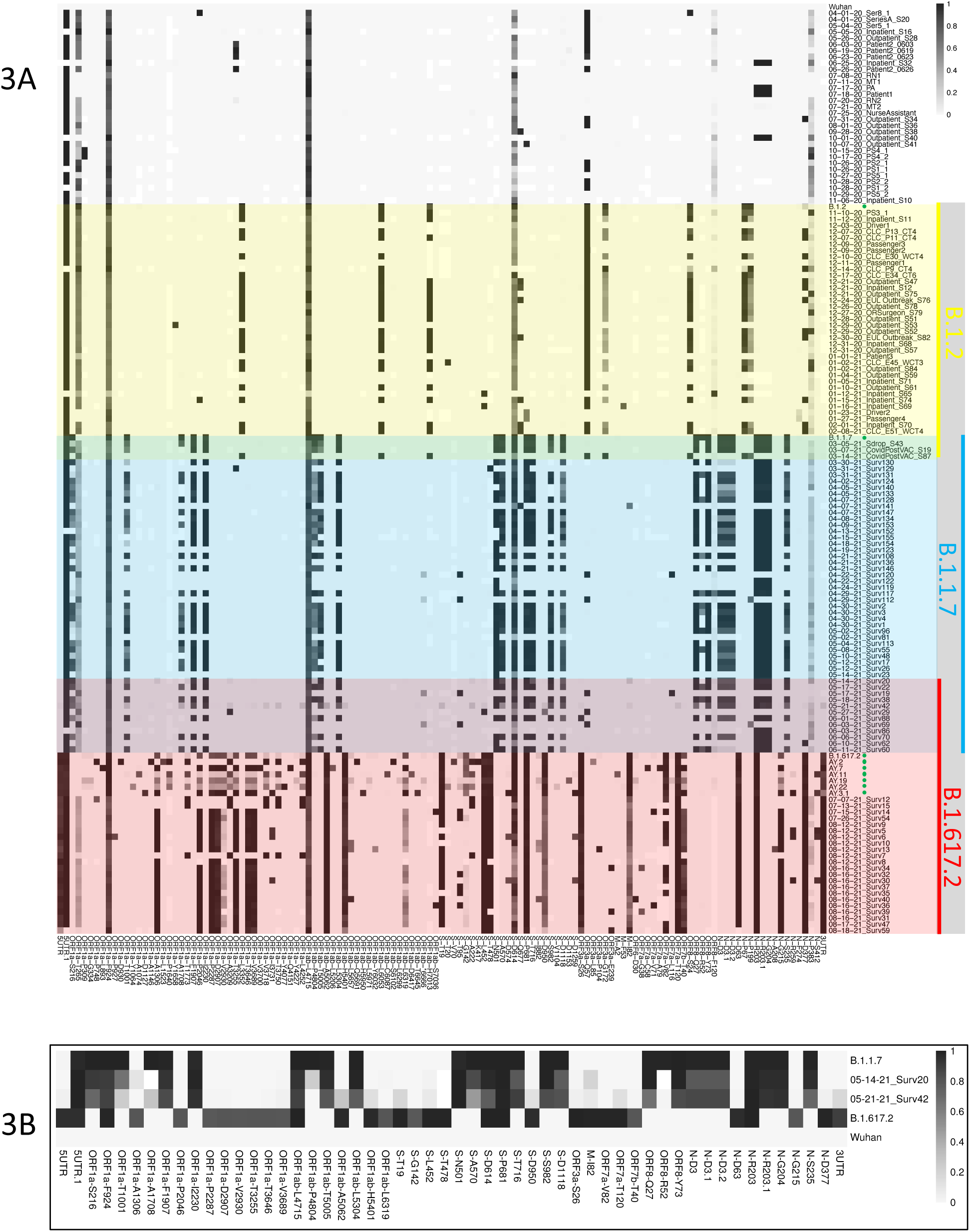
SARS-CoV-2 Sequence Complexity Integrated into Infection and Transmission Epidemiology. **a** Zebra-plot alignment of 140 sequences from the Cleveland VA study population, collected from April 2020 to August 2021 at 200 lineage defining signature positions. The grayscale bar (upper right) indicates the proportion of the alternate allele at each biallelic variant position. Integrated into the alignment of individual patient infection sequences and designated by green arrows are the Wuhan reference sequence (NC_045512.2), and aggregate plots for B.1.2 (51,069 sequences), B.1.1.7 (479,770 sequences), B.1.617.2 (258,910 sequences), AY2 (1,938 sequences), AY7 (3,443 sequences), AY11 (785 sequences), AY19 (106 sequences), AY22 (70 sequences), AY3.1 (5,363 sequences). Lineage defining signature positions are defined as a position in which >50% of samples from that lineage exhibits the alternate allele. **b** Sequence alignment and AAF proportion assessment of a SARS-CoV-2 infections Surv20 (May 14, 2021) and Surv42 (May 21, 2021) showing evidence of a B.1.1.7(Alpha)/B.1.617.2(Delta) infection.

Our observations of biallelic variant positions are consistent with those from other studies describing subconsensus SNPs or iSNVs [30–38]. To determine how often biallelic variations were included in SARS-CoV-2 sequences submitted to international repositories, we randomly selected and analyzed 110 additional COVID-19 infections (**Supplemental Table 4**) reported from 18 different laboratories or consortia with raw sequence data available in the sequence read archive (SRA) [39]; the same ≥80% coverage, ≥100X read-depth inclusion threshold was applied. Results summarized in **Supplemental Fig. 1** confirmed that all SRA files from the 110 external infections showed evidence of multiple biallelic iSNVs that would reflect the complexity of the SARS-CoV-2 infection (nucleotide positions characterized by biallelic iSNVs in our VA study population and the samples queried in the SAR are designated by a green dot in **Fig. 2** and **Supplemental Fig. 1**). Similar to observations from the VA Northeast Ohio Healthcare System patient population, 46 alternate biallelic iSNVs from a total of 87 variable positions did not clear the 50% consensus threshold. Among the 41 SNPs where AAFs did rise above the 50% threshold, the alternate allele would be reported in 668 occurrences and not reported in 312. Following the consensus sequence reporting convention across the 87 positions, 36% of the SARS-CoV-2 variations with AAFs between 5-50% (369 iSNVs) would not have been reported across the infections studied. Given that all SRA data queried showed this complexity, we asked whether attempts to report biallelic sequence variation were being made through the use of standard ambiguity codes [40] used to identify heterozygosity in diploid genetic systems. We first observed that the international data repositories understandably discourage the use of ambiguity codes given the substantial effort in attempting to curate sequence and determine lineage relationships as the pandemic continues to emerge. When we queried the database of 1,801,208 SARS-Cov-2 sequences accessible through Nextstrain/GISAID (available up to October 3, 2021), while the repository reports extensive nucleotide substitutions, results showed that 85.9% reported no ambiguity sequence codes suggesting no biallelic polymorphism in their SARS-CoV-2 sequences; 0.45% reported >10 suggesting potential for a complex multi-strain infection (**Supplemental Tables S6**). Given our presentation that most, if not all, SARS-CoV-2 full genome sequences are characterized by the occurrence of multiple biallelic positions, we have integrated appropriate ambiguity codes at positions where the AAF is >5% and observed that sequences are characterized by a minimum of 12 and maximum of 132 biallelic ambiguities. In head-to-head comparisons, nucleotide-specific substitution was higher in Nextstrain/GISAID compared to our study (average substitution rate = 1.69, st. dev. = 0.52), while biallelic ambiguity code usage is 100-fold lower (average substitution rate = 0.0097, st. dev. = 0.018); overall identification of mutations and ambiguity usage is summarized in **Supplemental Tables 5c to 5f**. Due to the increasing levels of SNVs observed over the time of sample collection, we report the average, minimum and maximum numbers of biallelic positions in quartiles (April 1 to August 1, 2020; September 28 to December 9, 2020; December 10, 2020 to April 13, 2021; April 15 to August 18, 2021), corresponding to breaks in sample availability (**Table 1**).

### Epidemiological Transitions of SARS-CoV-2 Infection Complexity

In addition to the accumulation of increased polymorphism across the viral genome over time, closer examination of the genomic variations in the summary zebra plot (so-called due to the gray-scale patterns; **Fig.3a**) revealed significant pattern shifts in the sequences that characterize the major variants in our sample population that correspond to epidemiological transition periods. Following the sample collection time series (**Fig. 3a**, April 2020 (top) to August 2021 (bottom)), four approximate variant signature time frames are observed - Pangolin lineages B.1 (no shading; April – November 2020), B.1.2 (yellow shading; November 2020 – March 2021), B.1.1.7 (Alpha; blue shading; March – June 2021) and B.1.617.2 (Delta; red shading; May – August 2021); green dots identify lineage-specific reference sequences compiled from frequencies of mutations deposited in GISAID for each lineage. Sequences within green and purple shades identify transition periods, B.1.2 to B.1.1.7 and B.1.1.7 to B.1.617.2, respectively. Of particular interest, the two exceptional samples identified in **Figs. 1d-1f** (samples Surv20 – May 14, 2021; Surv42 – May 21, 2021) are found in the B.1.1.7 to B.1.617.2 transition period. In evaluation of their genomic sequence data, variants are observed with an AAFs > 5% but not reaching major consensus, that are consistent with B.1.617.2-specific SNPs (**Fig 3b**). Following conventional practices, the major variants (with frequencies above 50% allele frequency cutoff) identified for these infections would have been reported as B.1.1.7 and not a mixed strain infection of B.1.1.7/B.1.617.2. Observation of these minor alleles above the 5% cutoff demonstrates early detection of the next major variant of concern (**Fig. 1f**).

A final summary of the distribution of SNVs observed in our study population is presented in **Fig. 4**. The data show that variation is significantly concentrated among the genes encoding structural proteins. This is particularly true for positions characterized by biallelic variation where AAF may not rise above 50% (Black components of stacked histogram bars); much of which is observed in the SARS-CoV-2 spike protein.

**Fig. 4.**
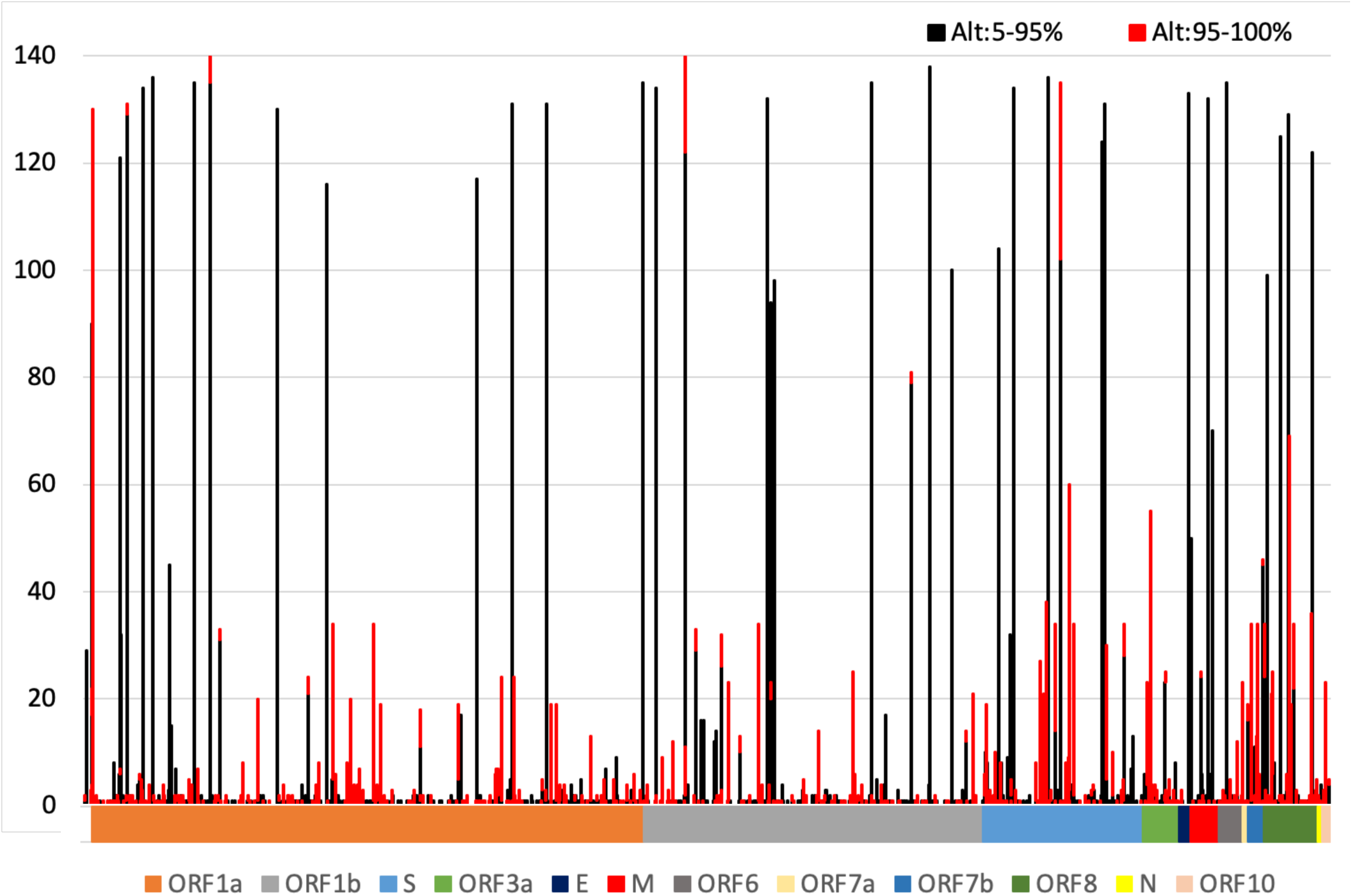
Summary of the Distribution of SARS-CoV-2 Sequence Variation and Heterogeneity. Counts of samples observed to show variation (among the 140 meeting the analytical threshold) across the SARS-CoV-2 genome. Black histogram components include those positions where biallelic variation was observed and may not have been included in reported consensus sequences. Red histogram components identify positions and occurrences where the alternate allele was observed at proportions >95% and would have been included in the reported consensus sequence. The multicolored bar below the graph marks genomic segments encoding different SARS-CoV-2 genes identified in the legend.

### Bottleneck Evaluations and Transmission Dynamics

Examination of the total infection sequence composition has also been considered in our study of infection transmission complexity from multiple exposure events. We performed bottleneck assessment using the exact beta-binomial method of Sobel-Leonard et al. [41], that recognizes AAF and RAF proportions instead of treating nucleotide sequence variation on a presence/absence basis taken by alternative models [42, 43]. These analyses focused on 11 donor-recipient pairs, some that have been studied previously [36–38]. While donors were known from transmission events that occurred during commuter van transport events, we performed reciprocal calculations (A to B and B to A) for all transmission pairs. Results from our overall analyses show the maximum likelihood estimate (MLE) of the bottlenecks among our transmission pairs was between 6 to 123 virions (**Fig. 5a**). These results include transmission events that occurred toward the end of pandemic Year-1 (December 2020 and January 2021) with a viral population characterized by greater genetic diversity compared to the first quarter of 2020 (**Fig. 1f**). Paired SNP proportions in the “transmission plots” (**Figs. 5b-5d**) show the AAF for each position in the SARS-CoV-2 genome between the respective infected donor (left) and subsequent recipient (right). In these graphs, most of the nucleotide-specific data are at or below the 5% alternate allele threshold and therefore would be designated the reference allele (>29,000 nucleotide positions). Results further indicate that among the majority of biallelic positions, the allele frequencies were transmitted in very similar proportions (AAF:RAF). As such, the average allele frequency differences (AFΔ) between donor and recipient pairs were minimal in all transmission scenarios, regardless of exposure details (0.09 to 0.17%; **Figures 5b** **to 5d**).

**Fig. 5.**
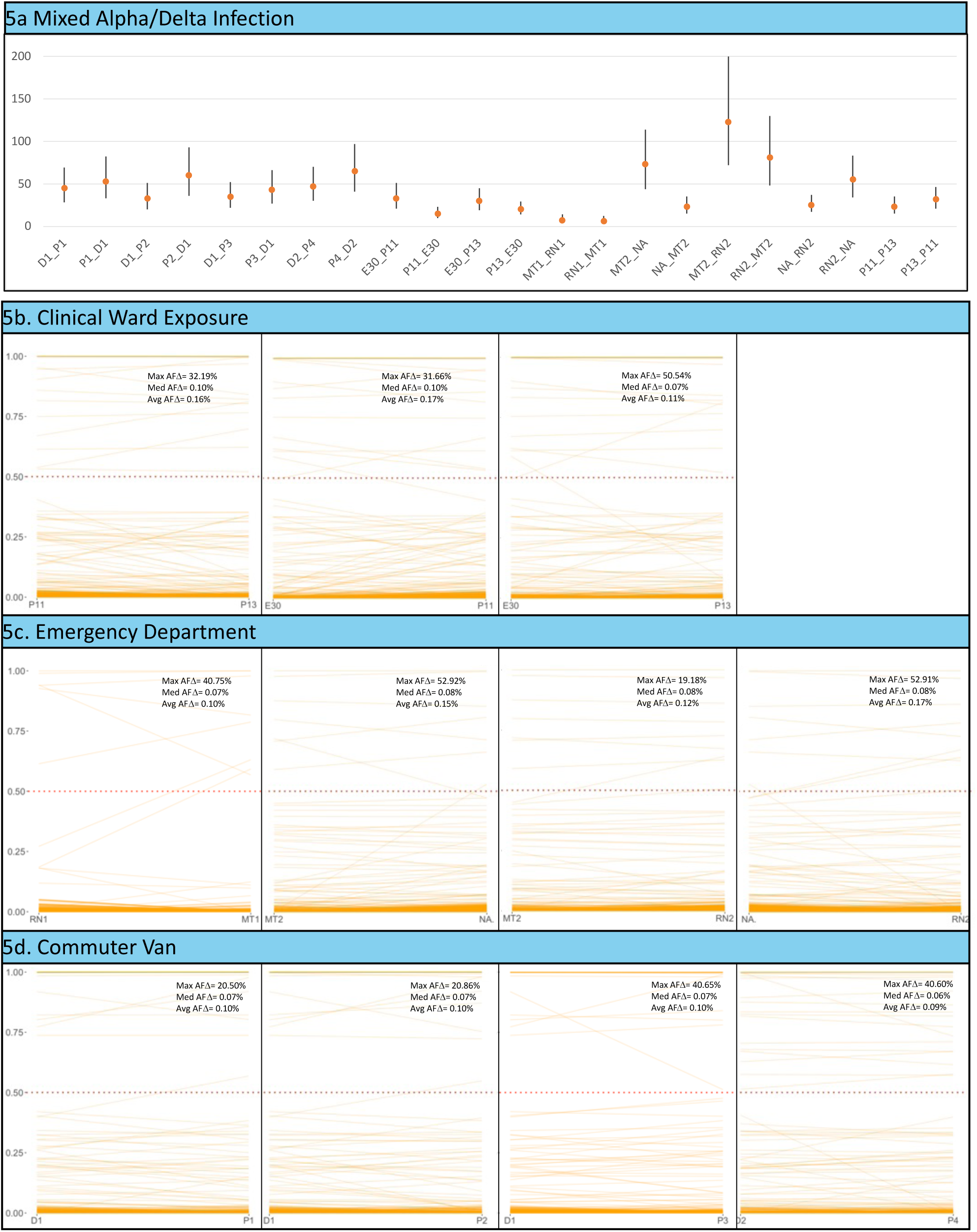
(following page). SARS-CoV-2 Transmission Bottleneck Assessment and Infection Sequence Complexity. **a** Summary of bottleneck evaluations for 11 donor-recipient pairs. Dates of transmission include June/July 2020 – MT1-RN1, MT2-RN2, NA-MT2, NA-RN2; November 2020/January 2021 – P11-P13, P11-E30, P13-E30; December 3, 2020 – D-P1, D1-P2, D1-P3; January 23, 2021 – D2-P4. Individual results provide the mean predicted number of virions for each transmission event and 95% CI. **b-d** Transmission graphs showing AAFs between the 11 infected donor and exposed recipient pairs. Infection complexity is demonstrated by an array of AAFs from 0.05 to 1.0 (y-axes). Transmission settings include a VA Clinical Ward b; a VA Emergency Department **c**; two independent commuter transport vans **d**. Inset date for each transmission graph show the maximum change in allele frequency (Max AFD), the median allele frequency change (Med AFD), and the average allele frequency change (Avg AFD). Allele sharing assessments were also performed for two randomly chosen pairs who did not work together or have known contact - CLCCB1 & EDCB6: Max: AFD = 99.95%, Med AFD = 0.081%, Avg AFD = 0.221%; SURVCB26 & EDCB5: Max AFD = 100.00%, Med AFD = 0.077%, Avg AFD = 0.249%.

### Association between Immunological Competence and SARS-CoV-2 Infection Complexity

Finally, given published reports of extensive SARS-CoV-2 sequence diversity in association with infected immunocompromised (IM-) individuals [44–47], we have evaluated the complexity of SARS-CoV-2 sequence in the context of the immunological conditions of the infected individuals in our study. Because the infections studied included asymptomatic individuals and those who receive primary care outside the VA Northeast Ohio Healthcare System, it was not possible to conduct a complete association study on this focus. Nevertheless, clinical evaluations of 16 individuals with sequence meeting our analysis inclusion threshold was possible. Given the observations shown in **Fig. 1f**, that genome-wide SNPs have increased during the pandemic, we compared SNPs detected in viruses from IM-with immunocompetent (IM+) individuals in relation to the time when their samples were collected. Results summarized in **Supplemental Fig. 2** showed that there were no obvious differences in the number of variations harbored by the virus from IM-compared to IM+ patients. Interestingly, the individuals identified to carry a B.1.1.7/B.1.617.2 mixed infection were discordant with respect to their immune status (one IM- the other IM+).

## DISCUSSION

Our study has revealed significant within host infection diversity of SARS-CoV-2 through full genome sequence analysis. We greatly appreciate the work of international data repositories such as GISAID and NextStrain for curating the millions of reported infections and believe that these repositories are a tremendous and invaluable asset for the monitoring of SARS-CoV-2. The shear amount of data that needs to be collected and annotated makes it understandable and perhaps necessary to represent these infections by a single consensus sequence. While the practice of reporting a single unambiguous sequence assists with tracking SARS-CoV-2 in regard to evolution and global distribution of lineages, emphasis on reporting the majority nucleotide signature significantly reduces our ability to detect emerging new variants that could ignite a new surge with public health impact. Additionally, the majority consensus approach limits our ability to understand the complexity of infections, dynamic changes in the viral population over time within infected individuals, and between donor and recipient individuals across transmission events.

In this study, we addressed the majority consensus problem as it relates to SARS-CoV-2 infection through the use of standard ambiguity codes [40]. All consensus sequences reported to GenBank from this study have included ambiguity codes. Taking this approach, our study accurately identified two individuals who harbored mixed B.1.1.7/B.1.617.2 infections in mid-May 2021. The collection of these samples preceded by one month the first reports from the State of Ohio that B.1.617.2 was present in the Cleveland area and occurred at a time when the City lifted its Civil Emergency Proclamation (May 28, 2021) and the State of Ohio rescinded its masking mandate and social distancing orders (June 2, 2021) [48]. Communicating knowledge of B.1.617.2 infection and transmission in Cleveland ahead of relaxing mitigation measures may have contributed to a more cautious public health strategy.

Additionally, nucleotides with biallelic mixtures observed repeatedly in multiple samples have included many of the functionally significant polymorphisms that have been reported in variants of concern. While widely distributed across the SARS-CoV-2 genome, biallelic mixtures are observed at amino acid positions within the SARS-CoV-2 spike protein’s receptor binding domain (e.g., L452R and T478K; B.1.617.2) that are prominent in lineage determination and associated with increased transmission efficiency (e.g., D614G). Moreover, there are numerous examples of alternate alleles observed in this study that did not clear the 50% threshold for inclusion in conventional consensus sequences that have subsequently become included as lineage defining SNPs in Omicron BA.1 (ORF1a:T3255I; S:T95I; S:K417N; S:H655Y; M:D3G; N:P13L; N:R203K; N:G204R) and/or BA.2 (ORF1a:T3255I; ORF6:D61L; S:T19I; S:G142D; S:K417N; S:H655Y; N:P13L; N:R203K; N:G204R) variants. These latter observations suggest that minority alleles in earlier SARS-CoV-2 infections may contribute to the continuing evolution of new variants of concern [49]. Confirming this potential will require consideration of minor alternate alleles as suggested here in more extensive whole genome analysis of SARS-CoV-2.

In our assessment of transmission from infected donors to recipients we were also interested to see that the overall sequence diversity was largely retained. This appeared to be true regardless of the AAF:RAF mixtures (**Figs. 5b** **to 5d**). These transmission plots provide both quantitative and qualitative perspectives on SARS-CoV-2 complexity. Many of the AAF:RAF mixtures were observed to be present in donor and recipient sequences at nearly identical ratios. For the events captured in commuter vans (two independent events) where the direction of transmission was known to be from infected drivers to uninfected passengers, this suggests that the majority of SARS-CoV-2 strain diversity is included during transmission and subsequent infection complexity [38]. This appeared to be true even though transmission occurred within the context of circulating air. Maximum likelihood estimates of the bottlenecks among our transmission pairs (van driver to passengers as well as others **Fig. 5a**) were similar or higher than those observed by Braun [35] and Lythgoe [34], but did not reach the highest values reported by Popa [32].

Finally, given reports that have suggested evolution of complexed populations of SARS-CoV-2 in immunocompromised people [44–47], we were interested to compare infections between IM+ and IM- individuals. While normalizing for overall complexity by comparing infections that occurred within time-similar cohorts, we did not observe increased numbers of iSNVs in IM- patients. This appears to differ from the observations of Al Khatib et al. who reported increased numbers of biallelic subconsensus variants (indicating AAF:RAF mixtures) in individuals classified as severe cases compared to individuals with mild cases [30] with comorbidities similar to the IM- patients studied here. It is also important to acknowledge that we were not able to study single patients over extended time series known to have extended for up to one year by other investigators [46]. Therefore, we did not have the opportunity to compare potential evolution of SARS-CoV-2 variants within individual IM+ and IM- patients over time.

The continuing dynamics of the COVID-19 pandemic have become increasingly complex and unpredictable. Underlying the challenge of COVID-19 transmission among humans, Sender et al. estimate that every infected person carries 1 billion to 100 billion virions during peak infection [50]. As a result, despite proof-reading activity within the SARS-CoV-2 RdRp that would limit mutation, efficient virus replication and transmission increasingly favors dispersal of mutations across abundant globally distributed infections (>460,000,000 to-date [22, 23]). As levels of immunity stimulated by infection and vaccination fluctuate in the human population ((a) no infection or vaccine exposure; (b) previous infection(s) and no vaccination; (c) no infections and vaccination(s); (d) previous infection(s) and vaccination(s)), the global SARS-CoV-2 population will encounter significant heterogeneity in acquired and natural human host defense mechanisms. Given the global disparities influencing access to, and uptake of vaccines, as well as availability and practice of non-pharmaceutical interventions, the virus has a free range for making thousands of deleterious as well as advantageous mutations. Therefore, significant opportunity exists for random chance to determine what future chapters of the pandemic will present.

This emphasizes the need to study genetic complexity of SARS-CoV-2 infections more intensively and requires increased examination to achieve a greater understanding of the capacity of this virus to evade our best efforts at mitigation. While there are efforts to sequence the virus and document how it is evolving, most of the scientific community recognizes that this effort needs to be increased [51–56]. Coupled with this effort, approaches to report characteristics of multi-strain complex infections must be considered to enable closer monitoring and full understanding of the virus’s evolutionary capacity.

Lest it be thought that under-reporting of the genetic diversity of infections occurs only with SARS-CoV-2 genome sequence data, consensus sequence reporting is the submission practice for most (if not all) infectious disease pathogens to national and international data repositories (https://VEuPathDB.org) from the US National Institute of Allergy and Infectious Diseases (NIAID) and the Wellcome Trust (example pathogens and vectors include biosecurity pathogens (e.g. *Yersinia pestis* (plague), Marburg virus, Ebola virus; pathogens of global concern [e.g. *Plasmodium*, *Mycobacterium tuberculosis*). Adequately addressing the under-reporting of infection complexity represented in data repositories has significant potential to alter vitally important characteristics of infectious diseases, including assessment of drug treatment efficacy and resistance and vaccine escape.

## METHODS

### Patient Enrollment and Informed Consent

As part of efforts to identify, provide care for infected individuals and limit the spread of COVID-19 within the VA Northeast Ohio Healthcare System, the Infection Control Department collected nasopharyngeal swab specimens from all individuals showing symptoms of a respiratory infection, all patients admitted to the hospital or undergoing invasive medical procedures, and asymptomatic close contacts of COVID-19 cases starting in March 2020 in accordance with Centers for Disease Control and Prevention (CDC) recommendations [57]. Since this time, we have analyzed over 300 samples from a variety of sources to investigate SARS-CoV-2 whole genome sequence variation. The study protocol was approved by the Cleveland VA Medical Center’s Institutional Review Board (IRB protocol number 1584025) with a waiver of consent.

### Patient Clinical Characteristics and Outcomes

Nasopharyngeal swab specimens were collected from patients and healthcare personnel. Clinical information was not available for healthcare personnel. For the patients, medical record review was performed to obtain information on age, medical conditions, medications, and severity of illness. COVID-19 was considered severe if oxygen saturation was <94% on room air or respiratory rate was >30 breaths/min or lung infiltrates >50% were present (https://www.covid19treatmentguidelines.nih.gov/overview/clinical-spectrum/). Patients were considered to be moderately or severely immunocompromised if they had conditions causing immunocompromise as defined by the Centers for Disease Control and Prevention (eg, receiving active cancer treatment for tumors or cancers of the blood, received an organ or stem cell transplant and taking medicine to suppress the immune system, advanced or untreated HIV infection, active treatment with high-dose corticosteroids or other drugs that suppress their immune response) (https://www.cdc.gov/coronavirus/2019-ncov/vaccines/recommendations/immuno.html).

### Patient Sampling and RNA Extraction

All nasopharyngeal swabs were immersed in 2 mL of universal transport medium (Copan Diagnostics, Murrieta, CA) and stored at -80°C. RNA was extracted from 140ul the stored sample to yield approximately 100ul of purified RNA following Qiagen protocols (either QIAamp Viral RNA Mini Kit or the QIAcube HT, automated 96-well plate format).

### Real-Time PCR Diagnosis of SARS-CoV-2 Infection

Real-time PCR (RT-PCR) amplification of RNA was performed following the Luna Universal One-Step RT-qPCR kit (New England Biolabs). A CFX 96 Touch Real-Time PCR Thermocycler/ Detection System (Bio-Rad) was used to quantify the amount of viral RNA present in each sample. The primer/probe set was designed as part of the Food and Drug Administration, Emergency Use Authorization protocol [58] was followed to amplify the SARS-CoV-2 nucleocapsid gene target sequence (Forward primer (2019-nCoV_N1-F) 5’- GACCCCAAAATCAGCGAAAT-3’; Reverse primer (2019-nCoV_N1-R) 5’- TCTGGTTACTGCCAGTTGAATCTG -3’; Probe (2019-nCoV_N1-P) design FAM-ACCCCGCATTACGTTTGGTGGACC-BHQ1). Temperature cycling conditions for the assay were as follows: 1 cycle of 55°C for 10 minutes (reverse transcription), 1 cycle of 95°C for 1 minute (initial denaturation), 45 cycles of 10 seconds at 95°C (denaturation) and 30 seconds at 60°C (extension). An in-plate fluorescence read was performed after each cycle of the 45-cycle amplification protocol. Melt curve analysis was added for verification at the end cycle. Standard curve calibration using know quantities of a SARS-CoV-2 RNA template (Exact Diagnostics)) were performed for each new primer/probe lot used in this assay. This standard contains RNA transcripts for the N gene of SARS-CoV-2 Additional controls for E, ORF1ab, RdRp, and S genes were available, but stability of N gene amplification did not prompt varying RT-PCR assessments. The target gene transcript was quantitated to 200,000 copies/mL using digital droplet PCR. Each RT-qPCR run contained positive and no template controls, in addition to the negative control from the extraction process.

### Whole-Genome Sequencing and Data Analysis

Whole-genome sequencing of SARS-CoV-2 was performed on 254 clinical samples from COVID-19 patients following library preparation reagents and protocols developed by Paragon Genomics (dual indexed CleanPlex SARS-CoV-2 Panel 918010 or 918011) [59]. Sequence was generated using an Illumina MiSeq sequencing platform. Raw sequences were aligned to COVID-19 reference sequence NC_045512.2. Primer sequences were trimmed from the ends of aligned reads using a masking file provided by Paragon Genomics using the software package fgbio. Samples were included in the final analysis if at least 80% of the genome was covered at > 100X read depth. Variants were called at positions with a minimum coverage of 100 reads with a base quality score >20. Positions were considered variable if any sample showed an alternate allele frequency at that position greater than 5%. These inclusion criteria are consistent and, in many ways, exceed benchmarks included in recently published quality assessments to evaluate efforts to sequence and describe SARS-CoV-2 genome sequence variation [60, 61].

We expanded our analyses to determine whether our observation of frequent biallelic iSNVs were characteristic of SARS-CoV-2 sequences submitted to international repositories. To perform this analysis, we randomly selected and analyzed 110 additional COVID-19 infections reported from 18 different laboratories or consortia with raw sequence data available in the sequence read archive (SRA) (**Supplemental Table 4**).

Inclusion of SARS-CoV-2 full genome sequence for all samples in the final analysis (generated by the current study or outside labs) required at least 80% of the genome was covered at > 100X read depth. Variants were called at positions with a minimum coverage of 100 reads with a base quality score >20 using Samtools mpileup v1.8. Positions were considered variable if any sample showed an alternate allele frequency at that position greater than 5%.

Consensus sequences were generated from this data by following conventional approaches, reporting the major nucleotide represented at each nucleotide position. Therefore, variant nucleotides in relation to NC_045512.2 were reported for any genomic position at which an alternate allele was detected at >50% in a sequenced sample. Additionally, sequences with ambiguity codes were also generated for allele frequencies between 5-95% using iVar Consensus (v.1.3) setting the minimum threshold t=0.95. Thus, in contrast to reporting sequence variation as limited to the standard nucleotide states (A-adenine; G-guanine; C-cytosine; U-uracil/T-thymine) iVar reports standard ambiguity codes to identify varying positions in the SARS-CoV-2 genome where more than one nucleotide state was observed at a frequency > 5% at a specific genome position (e.g. R= observation of A and G [40]; see Supplement). These consensus sequences inclusive of biallelic ambiguity codes have been reported to GenBank (NCBI – GenBank Accession numbers OM988245-OM988384; **Supplemental Table 3**) and GISAID. Additionally, raw sequencing data have been submitted to the SRA.

Evaluation of ambiguity code usage of sequence data available through Nextstrain was performed using *awk* on metadata available for 1,801,208 SARS-Cov-2 sequences accessible through Nextstrain up to October 3, 2021; data accessed on October 10, 2021.

## ACKNOWLEDGEMENTS

We thank the patients, administrators, engineering personnel, scientific and medical staff for their assistance with this study. We acknowledge Simone Edelheit for performing Illumina MiSeq sequencing and thank the staff of the Genomics Core Lab, Case Western Reserve University. We thank Niyati Thosani, Lily Liu and Rounak Fiegelman of Paragon Genomics for their technical assistance in optimizing SARS-CoV-sequencing library preparation and bioinformatic data analysis.

## AUTHOR CONTRIBUTIONS

**Conceptualization:** Ernest R. Chan, Lucas D. Jones, Marlin Linger, Jeffrey D. Kovach, Maria M. Torres-Teran, Audric Wertz, Curtis J. Donskey and Peter A. Zimmerman

**Data Curation:** Ernest R. Chan, Lucas D. Jones, Marlin Linger, Jeffrey D. Kovach, Maria M. Torres-Teran, Audric Wertz

**Formal Analysis:** Ernest R. Chan, Curtis J. Donskey and Peter A. Zimmerman

**Funding Acquisition:** Curtis J. Donskey and Peter A. Zimmerman

**Investigation:** Ernest R. Chan, Lucas D. Jones, Marlin Linger, Jeffrey D. Kovach, Maria M. Torres-Teran, Audric Wertz, Curtis J. Donskey and Peter A. Zimmerman

**Methodology:** Ernest R. Chan, Lucas D. Jones, Marlin Linger, Jeffrey D. Kovach, Maria M. Torres-Teran, Audric Wertz, Curtis J. Donskey and Peter A. Zimmerman

**Project Administration:** Ernest R. Chan, Curtis J. Donskey and Peter A. Zimmerman

**Resources**: Lucas D. Jones and Curtis J. Donskey

**Software:** Ernest R. Chan

**Supervision:** Curtis J. Donskey and Peter A. Zimmerman

**Validation:** Ernest R. Chan, Curtis J. Donskey and Peter A. Zimmerman

**Visualization:** Ernest R. Chan, Curtis J. Donskey and Peter A. Zimmerman

**Writing – Original Draft Preparation:** Ernest R. Chan, Peter A. Zimmerman

**Writing – Review & Editing:** Ernest R. Chan, Lucas D. Jones, Marlin Linger, Jeffrey D. Kovach, Maria M. Torres-Teran, Audric Wertz, Curtis J. Donskey and Peter A. Zimmerman

## Financial support

This work was supported by a Merit Review grant (CX001848) from the Department of Veterans Affairs to C.J.D and by a grant from the Department of Veterans Affairs Office of Research and Development as part of funding for VASeqCURE (grant number N/A) which in turn received funding from the American Rescue Plan Act funds..

## CONFLICTS OF INTEREST

C.J.D has received research grants from Clorox, Pfizer, and PDI. All other authors report no conflicts of interest relevant to this article.

## Supplemental Information

### TABLES

**Table S1. SARS-CoV-2 Gene-Specific SNV Counts**

**Table S2. Nucleotide Positions of SARS-CoV-2 SNV Detected in Current Study Population**

**Table S3. GenBank Accession Numbers for 140 Infection Consensus Sequences (Includes Biallelic Ambiguity Codes)**

**Table S4. SRA Identifier, Submitting Laboratory/Consortium and Sample Collection Date for 110 Infections**

**Table S5A. Count of Nucleotide Substitutions and Ambiguity Code Use in GISAID**

**Table S5B. Count of Nucleotide Substitutions and Ambiguity Code Use in Current Study**

**Table S5C. Table S5C. Frequency of Nucleotide Substitutions and Ambiguity Code Use in GISAID**

**Table S5D. Frequency of Nucleotide Substitutions and Ambiguity Code Use in Current Study**

**Table S5E. Average Difference in Substitutions Nextstrain/GISAID Compared to Current VA Study**

**Table S5F. Average Difference in Ambiguity Code Usage Nextstrain/GISAID Compared to Current VA Study**

**Table S6. Count of Nucleotide Substitutions and Ambiguity Code Use in Current Study**

### FIGURES

**Supplemental Figure 1.**
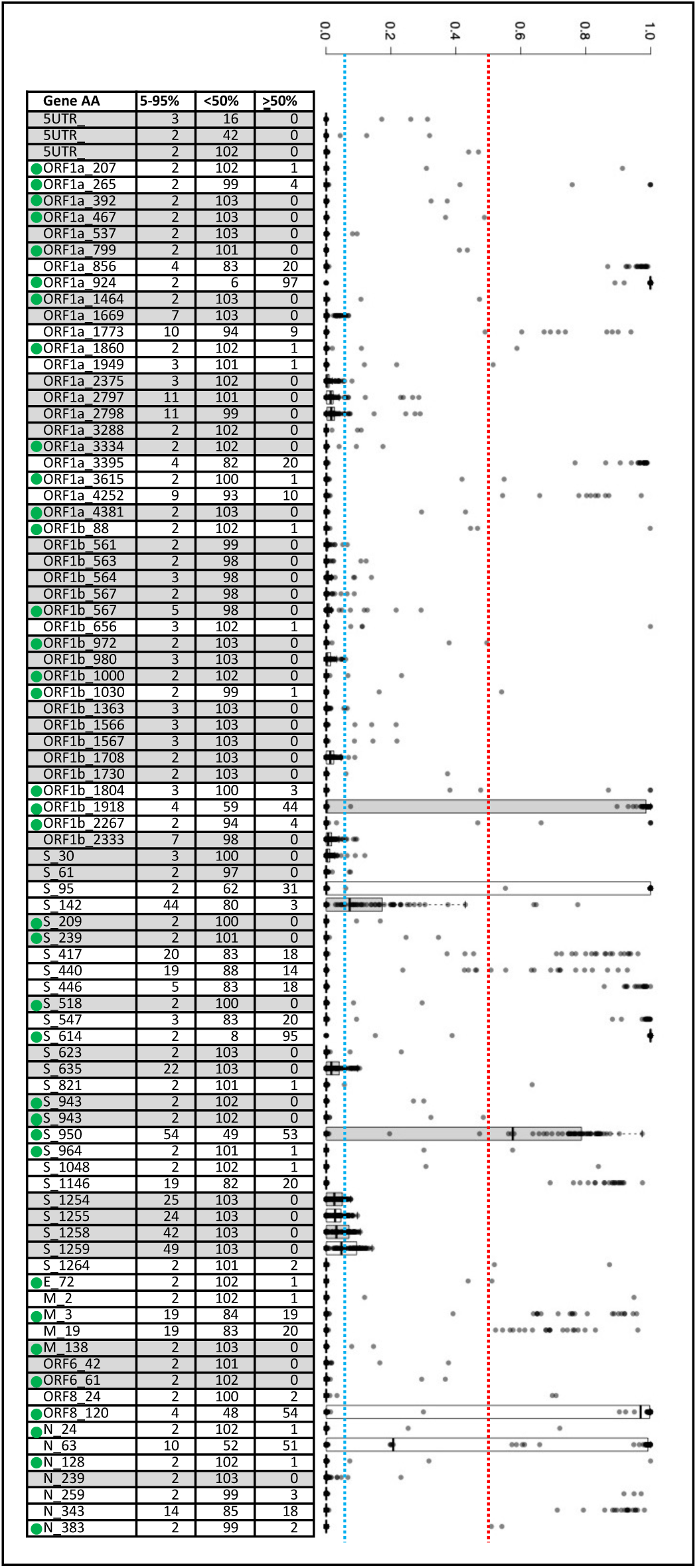
Assessment of Alternate and Reference Allele proportions (AAF:RAF) Across 110 Infections Accessed through the Sequence Read Archive (SRA). SRA data were accessed from 110 infections reported from 18 different laboratories or consortia as Illumina raw fastq files. These sources included AFRIMS, ATCC, Centre for DNA Fingerprinting and Diagnostics, COG-UK, CZB Covidtracker, Department of Medical Research, Disease Prevention and Control, Federal State Budgetary Scientific Institution Institute Of Experimental Medicine, Gonoshasthaya-RNA Molecular Diagnostics and Research Center, Igenbio Inc, Loma Linda University, National Public Health Laboratory of Malaysia, Network for Genomic Surveillance in South Africa (NGS_SA), The Hashemite University, University of California San Diego Microbiome Init, University of California Irvine, University of Montana, University of Wisconsin – Madison. Box and whisker plots illustrate the distribution of alternate allele frequencies for each mutation (median (bar); 25th and 75th percentiles (bottom and top of box, respectively); 10th and 90th percentiles (bottom and top whiskers, respectively)). The red dashed line at 50% identifies the demarcation at which the alternate allele is recognized and reported as the major consensus nucleotide that characterizes the infecting virus. These 87 mutations were chosen for further analysis because they all showed evidence of biallelic states at greater than the 5% background threshold and were observed in more than 2 infections (blue dashed line). Among these varying positions, 46 mutations did not reach an alternate allele frequency >50% for any of the 110 infections; 41 positions exceeded the 50% threshold for a portion of the 110 infections and would therefore have contributed to characterization of the infecting viral strain. The accompanying table enumerates the number of samples with AAF between 5 and 95%, <50% and >50%. Overall, among the 1,080 alternate alleles detected at a frequency >5%, just 657 (60.8%) were observed at an alternate allele frequency >50% necessary to be reported in consensus sequence. Nucleotide positions characterized by biallelic iSNVs in our VA study population (Fig. 2) and the samples queried in the SAR are designated by a green dot.

**Supplemental Figure 2.**
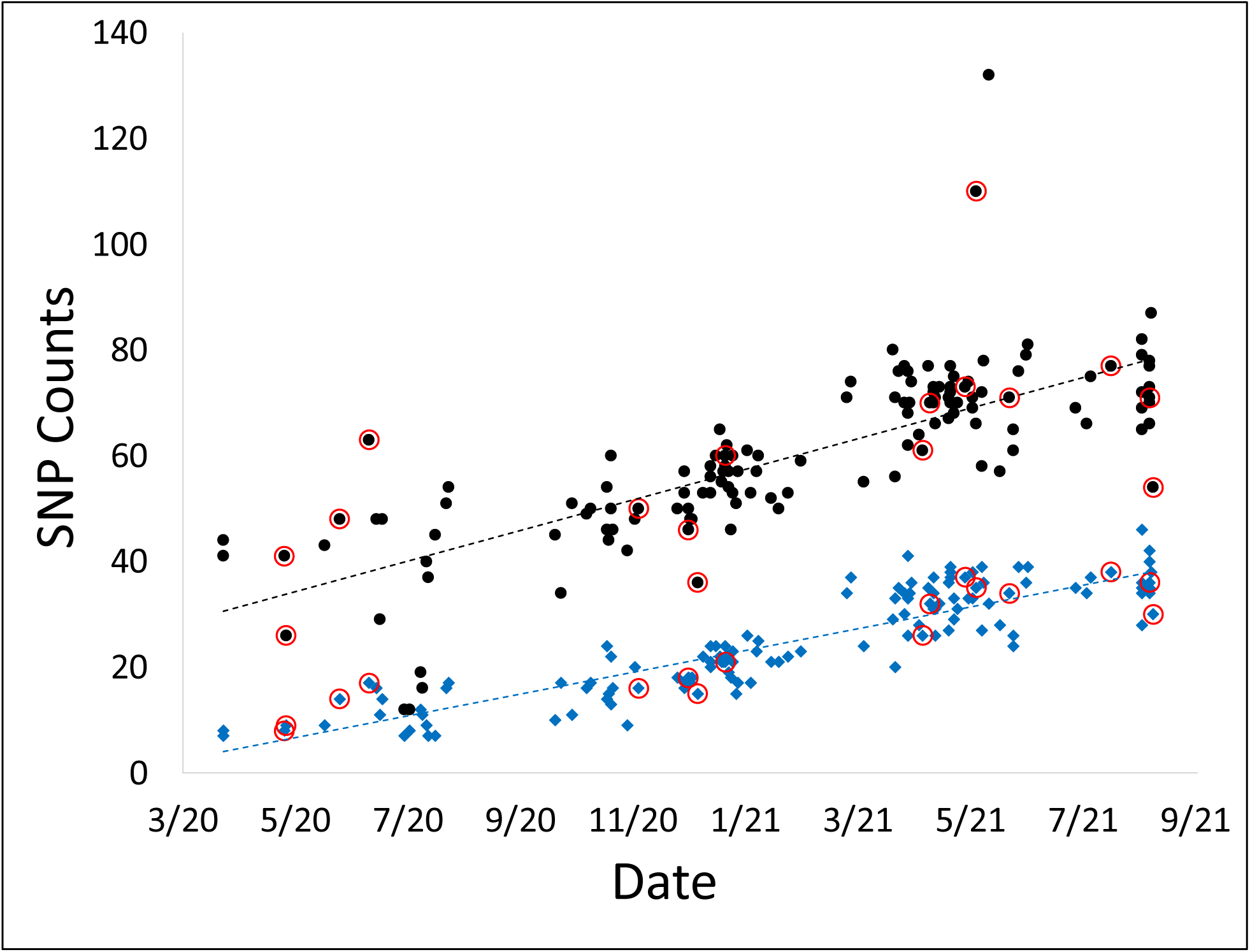
Comparison of SNVs between Immunocompetent (IM+) and Immunocompromised (IM-) Patients. IM- individuals are identified by red circles. Comparisons suggest that there is no difference in the SNVs observed at either threshold. Correlation between the number of alternate alleles identified in each sample and the sample collection date - AAF >5% (black circles), or AAF >50% (blue diamonds); samples were the 140 that were >80% 100x. Significant correlation was observed between an increase in AAFs during the sample collection time period (AAF >5%, Pearson R = 0.76112; AAF >50%, Pearson R = 0.89253; both P-values <0.0001).

## Notes

### Competing Interest Statement

CJD has received research grants from Clorox, Pfizer, and PDI. All other authors report no conflicts of interest relevant to this article.

## REFERENCES

1. Wu F, Zhao S, Yu B, Chen YM, Wang W, Song ZG, et al. A new coronavirus associated with human respiratory disease in China. Nature. 2020;579(7798):265–9. Epub 2020/02/06. doi: 10.1038/s41586-020-2008-3. PubMed PMID: 32015508; PubMed Central PMCID: PMCPMC7094943.

2. A novel coronavirus associated with a respiratory disease in Wuhan of Hubei province, China [Internet]. January 5, 2020. Available from: https://www.ncbi.nlm.nih.gov/nuccore/1798174254.

3. Marra MA, Jones SJ, Astell CR, Holt RA, Brooks-Wilson A, Butterfield YS, et al. The Genome sequence of the SARS-associated coronavirus. Science. 2003;300(5624):1399–404. Epub 2003/05/06. doi: 10.1126/science.1085953. PubMed PMID: 12730501; PubMed Central PMCID: PMCPMID: 12730501.

4. Rota PA, Oberste MS, Monroe SS, Nix WA, Campagnoli R, Icenogle JP, et al. Characterization of a novel coronavirus associated with severe acute respiratory syndrome. Science. 2003;300(5624):1394–9. Epub 2003/05/06. doi: 10.1126/science.1085952. PubMed PMID: 12730500; PubMed Central PMCID: PMCPMID: 12730500.

5. Zhu N, Zhang D, Wang W, Li X, Yang B, Song J, et al. A Novel Coronavirus from Patients with Pneumonia in China, 2019. N Engl J Med. 2020. Epub 2020/01/25. doi: 10.1056/NEJMoa2001017. PubMed PMID: 31978945.

6. van Dorp L, Richard D, Tan CCS, Shaw LP, Acman M, Balloux F. No evidence for increased transmissibility from recurrent mutations in SARS-CoV-2. Nature communications. 2020;11(1):5986. Epub 2020/11/27. doi: 10.1038/s41467-020-19818-2. PubMed PMID: 33239633; PubMed Central PMCID: PMCPMC7688939.

7. Maxmen A. One million coronavirus sequences: popular genome site hits mega milestone. Nature. 2021;593(7857):21. Epub 2021/04/25. doi: 10.1038/d41586-021-01069-w. PubMed PMID: 33893460.

8. Duchene S, Featherstone L, Haritopoulou-Sinanidou M, Rambaut A, Lemey P, Baele G. Temporal signal and the phylodynamic threshold of SARS-CoV-2. Virus Evol. 2020;6(2):veaa061. Epub 20200819. doi: 10.1093/ve/veaa061. PubMed PMID: 33235813; PubMed Central PMCID: PMCPMC7454936.

9. Rambaut A, Holmes EC, O’Toole A, Hill V, McCrone JT, Ruis C, et al. A dynamic nomenclature proposal for SARS-CoV-2 lineages to assist genomic epidemiology. Nat Microbiol. 2020;5(11):1403–7. Epub 2020/07/17. doi: 10.1038/s41564-020-0770-5. PubMed PMID: 32669681.

10. O’Toole A, Scher E, Underwood A, Jackson B, Hill V, McCrone JT, et al. Assignment of epidemiological lineages in an emerging pandemic using the pangolin tool. Virus Evol. 2021;7(2):veab064. Epub 20210730. doi: 10.1093/ve/veab064. PubMed PMID: 34527285; PubMed Central PMCID: PMCPMC8344591.

11. Konings F, Perkins MD, Kuhn JH, Pallen MJ, Alm EJ, Archer BN, et al. SARS-CoV-2 Variants of Interest and Concern naming scheme conducive for global discourse. Nat Microbiol. 2021;6(7):821–3. doi: 10.1038/s41564-021-00932-w. PubMed PMID: 34108654.

12. World Health Organization. W.H.O. announces simple, easy-to-say labels for SARS-CoV-2 variants of interest and concern - 31 May 2021. World Health Organization; 2021.

13. Denison MR, Graham RL, Donaldson EF, Eckerle LD, Baric RS. Coronaviruses: an RNA proofreading machine regulates replication fidelity and diversity. RNA Biol. 2011;8(2):270–9. Epub 2011/05/20. doi: 10.4161/rna.8.2.15013. PubMed PMID: 21593585; PubMed Central PMCID: PMCPMC3127101.

14. Minskaia E, Hertzig T, Gorbalenya AE, Campanacci V, Cambillau C, Canard B, et al. Discovery of an RNA virus 3’->5’ exoribonuclease that is critically involved in coronavirus RNA synthesis. Proc Natl Acad Sci U S A. 2006;103(13):5108–13. Epub 2006/03/22. doi: 10.1073/pnas.0508200103. PubMed PMID: 16549795; PubMed Central PMCID: PMCPMC1458802.

15. Duffy S, Shackelton LA, Holmes EC. Rates of evolutionary change in viruses: patterns and determinants. Nat Rev Genet. 2008;9(4):267–76. Epub 2008/03/06. doi: 10.1038/nrg2323. PubMed PMID: 18319742.

16. Rausch JW, Capoferri AA, Katusiime MG, Patro SC, Kearney MF. Low genetic diversity may be an Achilles heel of SARS-CoV-2. Proc Natl Acad Sci U S A. 2020;117(40):24614–6. Epub 2020/09/23. doi: 10.1073/pnas.2017726117. PubMed PMID: 32958678; PubMed Central PMCID: PMCPMC7547206.

17. Chaw SM, Tai JH, Chen SL, Hsieh CH, Chang SY, Yeh SH, et al. The origin and underlying driving forces of the SARS-CoV-2 outbreak. J Biomed Sci. 2020;27(1):73. Epub 20200607. doi: 10.1186/s12929-020-00665-8. PubMed PMID: 32507105; PubMed Central PMCID: PMCPMC7276232.

18. Li X, Zai J, Zhao Q, Nie Q, Li Y, Foley BT, et al. Evolutionary history, potential intermediate animal host, and cross-species analyses of SARS-CoV-2. J Med Virol. 2020;92(6):602–11. Epub 20200311. doi: 10.1002/jmv.25731. PubMed PMID: 32104911; PubMed Central PMCID: PMCPMC7228310.

19. Zhao Z, Li H, Wu X, Zhong Y, Zhang K, Zhang YP, et al. Moderate mutation rate in the SARS coronavirus genome and its implications. BMC Evol Biol. 2004;4:21. Epub 2004/06/30. doi: 10.1186/1471-2148-4-21. PubMed PMID: 15222897; PubMed Central PMCID: PMCPMC446188.

20. Khare S, Gurry C, Freitas L, Schultz MB, Bach G, Diallo A, et al. GISAID’s Role in Pandemic Response. China CDC Weekly. 2021;3(49):1049–51.

21. Hadfield J, Megill C, Bell SM, Huddleston J, Potter B, Callender C, et al. Nextstrain: real-time tracking of pathogen evolution. Bioinformatics. 2018;34(23):4121–3. doi: 10.1093/bioinformatics/bty407. PubMed PMID: 29790939; PubMed Central PMCID: PMCPMC6247931.

22. Dong E, Du H, Gardner L. An interactive web-based dashboard to track COVID-19 in real time. Lancet Infect Dis. 2020;20(5):533–4. Epub 2020/02/23. doi: 10.1016/S1473-3099(20)30120-1. PubMed PMID: 32087114; PubMed Central PMCID: PMCPMC7159018.

23. Bosch J, Wilson A, O’’Neil K, Zimmerman PA. COVID-19Predict - Predicting Pandemic Trends [Dashboard]. medRxiv2020 [updated December 29, 2020December 29, 2020]. Available from: https://covid19predict.com.

24. Mercatelli D, Giorgi FM. Geographic and Genomic Distribution of SARS-CoV-2 Mutations. Front Microbiol. 2020;11:1800. Epub 2020/08/15. doi: 10.3389/fmicb.2020.01800. PubMed PMID: 32793182; PubMed Central PMCID: PMCPMC7387429.

25. Puenpa J, Suwannakarn K, Chansaenroj J, Nilyanimit P, Yorsaeng R, Auphimai C, et al. Molecular epidemiology of the first wave of severe acute respiratory syndrome coronavirus 2 infection in Thailand in 2020. Scientific reports. 2020;10(1):16602. Epub 2020/10/08. doi: 10.1038/s41598-020-73554-7. PubMed PMID: 33024144; PubMed Central PMCID: PMCPMC7538975.

26. Yang HC, Chen CH, Wang JH, Liao HC, Yang CT, Chen CW, et al. Analysis of genomic distributions of SARS-CoV-2 reveals a dominant strain type with strong allelic associations. Proc Natl Acad Sci U S A. 2020;117(48):30679–86. Epub 2020/11/14. doi: 10.1073/pnas.2007840117. PubMed PMID: 33184173; PubMed Central PMCID: PMCPMC7720151.

27. Yuan F, Wang L, Fang Y, Wang L. Global SNP analysis of 11,183 SARS-CoV-2 strains reveals high genetic diversity. Transbound Emerg Dis. 2020. Epub 2020/11/19. doi: 10.1111/tbed.13931. PubMed PMID: 33207070; PubMed Central PMCID: PMCPMC7753349.

28. Centers for Disease Control and Prevention. Guidance for Reporting SARS-CoV-2 Sequencing Results Atlanta, GA2021 [August 8, 2021]. Available from: https://www.cdc.gov/coronavirus/2019-ncov/lab/resources/reporting-sequencing-guidance.html.

29. Global Initiative on Sharing All Influenza Data (GISAID). Submitting Data to the EpiFlu™ Database 2021 [August 8, 2021]. Available from: https://www.gisaid.org/epiflu-applications/submitting-data-to-epiflutm/.

30. Al Khatib HA, Benslimane FM, Elbashir IE, Coyle PV, Al Maslamani MA, Al-Khal A, et al. Within-Host Diversity of SARS-CoV-2 in COVID-19 Patients With Variable Disease Severities. Front Cell Infect Microbiol. 2020;10:575613. Epub 2020/10/31. doi: 10.3389/fcimb.2020.575613. PubMed PMID: 33123498; PubMed Central PMCID: PMCPMC7572854.

31. Du P, Ding N, Li J, Zhang F, Wang Q, Chen Z, et al. Genomic surveillance of COVID-19 cases in Beijing. Nature communications. 2020;11(1):5503. Epub 2020/11/01. doi: 10.1038/s41467-020-19345-0. PubMed PMID: 33127911; PubMed Central PMCID: PMCPMC7603498.

32. Popa A, Genger JW, Nicholson MD, Penz T, Schmid D, Aberle SW, et al. Genomic epidemiology of superspreading events in Austria reveals mutational dynamics and transmission properties of SARS-CoV-2. Sci Transl Med. 2020;12(573). Epub 2020/11/25. doi: 10.1126/scitranslmed.abe2555. PubMed PMID: 33229462; PubMed Central PMCID: PMCPMC7857414.

33. Wang Y, Wang D, Zhang L, Sun W, Zhang Z, Chen W, et al. Intra-host variation and evolutionary dynamics of SARS-CoV-2 populations in COVID-19 patients. Genome Med. 2021;13(1):30. Epub 2021/02/24. doi: 10.1186/s13073-021-00847-5. PubMed PMID: 33618765; PubMed Central PMCID: PMCPMC7898256.

34. Lythgoe KA, Hall M, Ferretti L, de Cesare M, MacIntyre-Cockett G, Trebes A, et al. SARS-CoV-2 within-host diversity and transmission. Science. 2021;372(6539). Epub 2021/03/11. doi: 10.1126/science.abg0821. PubMed PMID: 33688063; PubMed Central PMCID: PMCPMC8128293.

35. Braun KM, Moreno GK, Wagner C, Accola MA, Rehrauer WM, Baker DA, et al. Acute SARS-CoV-2 infections harbor limited within-host diversity and transmit via tight transmission bottlenecks. PLoS pathogens. 2021;17(8):e1009849. Epub 20210823. doi: 10.1371/journal.ppat.1009849. PubMed PMID: 34424945; PubMed Central PMCID: PMCPMC8412271.

36. Chan ER, Jones LD, Redmond SN, Navas ME, Kachaluba NM, Zabarsky TF, et al. Use of whole-genome sequencing to investigate a cluster of severe acute respiratory syndrome coronavirus 2 (SARS-CoV-2) infections in emergency department personnel. Infect Control Hosp Epidemiol. 2021:1–3. Epub 2021/05/05. doi: 10.1017/ice.2021.208. PubMed PMID: 33941299; PubMed Central PMCID: PMCPMC8144813.

37. Jones LD, Chan ER, Cadnum JL, Redmond SN, Navas ME, Zabarsky TF, et al. Investigation of a cluster of severe acute respiratory syndrome coronavirus 2 (SARS-CoV-2) infections in a hospital administration building. Infect Control Hosp Epidemiol. 2022:1–7. Epub 20220222. doi: 10.1017/ice.2022.45. PubMed PMID: 35189996.

38. Jones LD, Chan ER, Zabarsky TF, Cadnum JL, Navas ME, Redmond SN, et al. Transmission of Severe Acute Respiratory Syndrome Coronavirus 2 (SARS-CoV-2) in a Patient Transport Van. Clin Infect Dis. 2022;74(2):339–42. doi: 10.1093/cid/ciab347. PubMed PMID: 33893474; PubMed Central PMCID: PMCPMC8135457.

39. Rigden DJ, Fernandez XM. The 2022 Nucleic Acids Research database issue and the online molecular biology database collection. Nucleic Acids Res. 2022;50(D1):D1–D10. doi: 10.1093/nar/gkab1195. PubMed PMID: 34986604; PubMed Central PMCID: PMCPMC8728296.

40. Cornish-Bowden A. Nomenclature for incompletely specified bases in nucleic acid sequences: recommendations 1984. Nucleic Acids Res. 1985;13(9):3021–30. doi: 10.1093/nar/13.9.3021. PubMed PMID: 2582368; PubMed Central PMCID: PMCPMC341218.

41. Sobel Leonard A, Weissman DB, Greenbaum B, Ghedin E, Koelle K. Transmission Bottleneck Size Estimation from Pathogen Deep-Sequencing Data, with an Application to Human Influenza A Virus. J Virol. 2017;91(14). Epub 20170626. doi: 10.1128/JVI.00171-17. PubMed PMID: 28468874; PubMed Central PMCID: PMCPMC5487570.

42. Emmett KJ, Lee A, Khiabanian H, Rabadan R. High-resolution Genomic Surveillance of 2014 Ebolavirus Using Shared Subclonal Variants. PLoS Curr. 2015;7. Epub 20150209. doi: 10.1371/currents.outbreaks.c7fd7946ba606c982668a96bcba43c90. PubMed PMID: 25737802; PubMed Central PMCID: PMCPMC4339230.

43. Poon LL, Song T, Rosenfeld R, Lin X, Rogers MB, Zhou B, et al. Quantifying influenza virus diversity and transmission in humans. Nat Genet. 2016;48(2):195–200. Epub 20160104. doi: 10.1038/ng.3479. PubMed PMID: 26727660; PubMed Central PMCID: PMCPMC4731279.

44. Bansal N, Raturi M, Bansal Y. SARS-CoV-2 variants in immunocompromised COVID-19 patients: The underlying causes and the way forward. Transfus Clin Biol. 2021. Epub 20211229. doi: 10.1016/j.tracli.2021.12.006. PubMed PMID: 34973463; PubMed Central PMCID: PMCPMC8714679.

45. Chen L, Zody MC, Di Germanio C, Martinelli R, Mediavilla JR, Cunningham MH, et al. Emergence of Multiple SARS-CoV-2 Antibody Escape Variants in an Immunocompromised Host Undergoing Convalescent Plasma Treatment. mSphere. 2021;6(4):e0048021. Epub 20210825. doi: 10.1128/mSphere.00480-21. PubMed PMID: 34431691; PubMed Central PMCID: PMCPMC8386433.

46. Nussenblatt V, Roder AE, Das S, de Wit E, Youn JH, Banakis S, et al. Year-long COVID-19 infection reveals within-host evolution of SARS-CoV-2 in a patient with B cell depletion. J Infect Dis. 2021. Epub 20211223. doi: 10.1093/infdis/jiab622. PubMed PMID: 34940844; PubMed Central PMCID: PMCPMC8755281.

47. Weigang S, Fuchs J, Zimmer G, Schnepf D, Kern L, Beer J, et al. Within-host evolution of SARS-CoV-2 in an immunosuppressed COVID-19 patient as a source of immune escape variants. Nature communications. 2021;12(1):6405. Epub 20211104. doi: 10.1038/s41467-021-26602-3. PubMed PMID: 34737266; PubMed Central PMCID: PMCPMC8568958.

48. Health CDoP. An Overview of COVID-19 in Cleveland, Ohio: 18-Monty Report (February 2020-August 2021). Cleveland, Ohio2021. p. 46.

49. Bansal K, Kumar S. Mutational cascade of SARS-CoV-2 leading to evolution and emergence of omicron variant. Virus Res. 2022;315:198765. Epub 20220331. doi: 10.1016/j.virusres.2022.198765. PubMed PMID: 35367284; PubMed Central PMCID: PMCPMC8968180.

50. Sender R, Bar-On YM, Gleizer S, Bernshtein B, Flamholz A, Phillips R, et al. The total number and mass of SARS-CoV-2 virions. Proc Natl Acad Sci U S A. 2021;118(25). doi: 10.1073/pnas.2024815118. PubMed PMID: 34083352; PubMed Central PMCID: PMCPMC8237675.

51. Boehm E, Kronig I, Neher RA, Eckerle I, Vetter P, Kaiser L, et al. Novel SARS-CoV-2 variants: the pandemics within the pandemic. Clinical microbiology and infection : the official publication of the European Society of Clinical Microbiology and Infectious Diseases. 2021;27(8):1109–17. Epub 20210517. doi: 10.1016/j.cmi.2021.05.022. PubMed PMID: 34015535; PubMed Central PMCID: PMCPMC8127517.

52. Callaway E. Multitude of coronavirus variants found in the US - but the threat is unclear. Nature. 2021;591(7849):190. doi: 10.1038/d41586-021-00564-4. PubMed PMID: 33674807.

53. Crawford DC, Williams SM. Global variation in sequencing impedes SARS-CoV-2 surveillance. PLoS Genet. 2021;17(7):e1009620. Epub 20210715. doi: 10.1371/journal.pgen.1009620. PubMed PMID: 34264957; PubMed Central PMCID: PMCPMC8282079.

54. Hodcroft EB, De Maio N, Lanfear R, MacCannell DR, Minh BQ, Schmidt HA, et al. Want to track pandemic variants faster? Fix the bioinformatics bottleneck. Nature. 2021;591(7848):30–3. doi: 10.1038/d41586-021-00525-x. PubMed PMID: 33649511.

55. Maxmen A. Omicron blindspots: why it’s hard to track coronavirus variants. Nature. 2021;600(7890):579. doi: 10.1038/d41586-021-03698-7. PubMed PMID: 34916668.

56. Maxmen A. Why US coronavirus tracking can’t keep up with concerning variants. Nature. 2021;592(7854):336–7. doi: 10.1038/d41586-021-00908-0. PubMed PMID: 33828280.

57. Centers for Disease Control and Prevention. Interim U.S. guidance for risk assessment and work restrictions for healthcare personnel 160 with potential exposure to COVID-19. Atlanta, GA2021 [updated February 25,2021August 12, 2021]. Available from: https://www.cdc.gov/coronavirus/2019-ncov/php/contact-tracing/contact-tracing-plan/contact-tracing.html.

58. United States Food and Drug Administration. Accelerated Emergency Use Authorization (EUA) Summary Covid-19 RT-PCR Test (Laboratory Corporation of America). Lab Corp; February 29, 2020.

59. Li C, Debruyne DN, Spencer J, Kapoor V, Liu LY, Zhou B, et al. Highly sensitive and full-genome interrogation of SARS-CoV-2 using multiplexed PCR enrichment followed by next-generation sequencing. biorRxiv. 2020. Epub May 18, 2020. doi: https://doi.org/10.1101/2020.03.12.988246.

60. Jacot D, Pillonel T, Greub G, Bertelli C. Assessment of SARS-CoV-2 Genome Sequencing: Quality Criteria and Low-Frequency Variants. J Clin Microbiol. 2021;59(10):e0094421. Epub 20210728. doi: 10.1128/JCM.00944-21. PubMed PMID: 34319802; PubMed Central PMCID: PMCPMC8451431.

61. Wegner F, Roloff T, Huber M, Cordey S, Ramette A, Gerth Y, et al. External Quality Assessment of SARS-CoV-2 Sequencing: an ESGMD-SSM Pilot Trial across 15 European Laboratories. J Clin Microbiol. 2022;60(1):e0169821. Epub 20211110. doi: 10.1128/JCM.01698-21. PubMed PMID: 34757834.

